# Dynamic PD-L1 Regulation Shapes Tumor Immune Escape and Response to Immunotherapy

**DOI:** 10.1101/2025.10.28.685116

**Authors:** Bruce Pell, Aigerim Kalizhanova, Aisha Tursynkozha, Denise Dengi, Ardak Kashkynbayev, Yang Kuang

## Abstract

A major challenge in cancer treatment is the ability of tumor cells to adapt to immunotherapy through immune escape, often mediated by the PD-1/PD-L1 pathway. To investigate this, we adapted an ordinary differential equation model of combination therapy, incorporating the dynamics of the immune checkpoint inhibitor Avelumab and the immunostimulant NHS-muIL12. Using literature-derived parameter values from a previous study, we refitted a single parameter across therapies, which showed that PD-L1 expression increased with immunotherapy, while Avelumab blocked its functional signaling, preventing PD-L1 from suppressing T-cell activity. Incorporating therapy-dependent, dynamically regulated PD-L1 expression enabled a biologically grounded mechanism to reproduce experimental observations, leading us to formulate PD-L1 tumor expression as a dynamic variable (*ϵ*) and providing a mechanistic basis for both therapeutic synergy and treatment failure. Our results indicate that tumor resistance is linked to dose-dependent upregulation of PD-L1 following NHS-muIL12 treatment, explaining treatment failure, while PD-1/PD-L1 blockade in combination therapy enables effective anti-tumor immune responses.

## 1. Introduction

Cancerous tumors are not static entities but dynamically evolve under immune and therapeutic pressure, often developing sophisticated strategies to evade immune surveillance [1–3]. Understanding these adaptive mechanisms, particularly the intricate regulation of immune checkpoints like Programmed Cell Death Protein 1 (PD-1) or its ligand (PD-L1), is important for overcoming resistance to modern immunotherapies and for generating personalized treatment plans for patients [4–6]. The PD-1/PD-L1 signaling axis functions as a regulator of tumor immunity by suppressing T-cell activation, proliferation, and cytotoxic capacity [7]. Activated T-cells express the PD-1 receptor, and when it binds to its ligand, PD-L1, it delivers an inhibitory signal that deactivates the T-cell. Many cancers have co-opted this natural mechanism to evade the immune system by overexpressing PD-L1 on their surface, essentially neutralizing attacking T-cells [8].

The development of immune checkpoint inhibitors, which are monoclonal antibodies that physically block the interaction between PD-1 and PD-L1, has revolutionized cancer therapy [4,8]. Avelumab is an immunotherapy drug that acts as an immune checkpoint inhibitor to treat specific types of advanced cancer, including Merkel cell carcinoma, urothelial cancer, and renal cell carcinoma [9,10]. By binding to PD-L1, Avelumab blocks its interaction with the PD-1 receptor on T-cells, thereby preventing the deactivation of the anti-tumor immune response and effectively allowing cytotoxic T-lymphocytes to kill cancer cells [11]. While effective, checkpoint blockade alone often fails due to insufficient immune activation within the tumor microenvironment. For this reason, combination strategies that pair checkpoint inhibition with immunostimulatory agents have gained interest. One such agent is NHS-muIL12, a tumor-targeting immunocytokine that combines a tumor-targeting antibody with the immune-stimulating cytokine interleukin-12 (IL-12) [12]. IL-12 is a potent cytokine that stimulates the proliferation and activation of T-cells and Natural Killer (NK) cells [13]. Preclinical studies demonstrated that NHS-muIL12 increases immune infiltration and activity, while Avelumab prevents tumor-mediated suppression, leading to synergistic anti-tumor effects [14]. See Figure 1 for a schematic illustration of the main dynamics.

**Figure 1.**
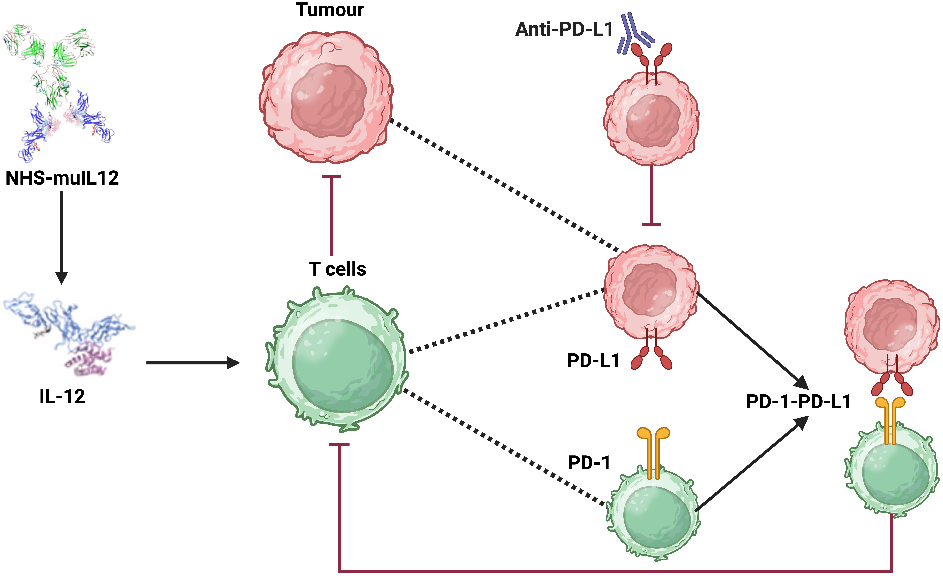
Schematic illustration of the synergistic anti-tumor mechanisms of NHS-muIL12 and anti-PD-L1 (Avelumab) checkpoint blockade within the tumor microenvironment. The diagram shows how NHS-muIL12 delivers IL-12 to promote T cell activation, while anti-PD-L1 antibody therapy overcomes tumor-induced immune suppression by disrupting the PD-1/PD-L1 axis. Created in BioRender. Tursynkozha, A. (2025) https://BioRender.com/4yxwllp.

Mathematical modeling has emerged as a powerful tool to provide novel insight into cancer biology, tumor growth, and treatment response [15–18]. A comprehensive coverage of these topics can be found in Kuang et al. [19].These quantitative frameworks are essential for deciphering the intricate, non-linear interactions between cancer cells, diverse immune populations, and therapeutic treatments, which are often challenging to isolate and measure experimentally [20,21]. Since mathematical models can be designed with specific mechanisms and pathways in mind, they provide a way to generate testable, data-driven hypotheses and optimize treatment strategies when parameterized to data [22–25]. For example, Meade et al. used a mathematical model of prostate cancer to develop novel indicators of treatment failure. Their work, based on an evolutionary perspective, led to the hypothesis that the ratio of androgen to prostate-specific antigen (PSA) could serve as a powerful prognostic biomarker for predicting resistance to therapy [24,25].

In a significant effort to understand the intricate dynamics governing tumor cell proliferation, immune responses, and the balance of therapeutic interventions, Nikolopoulou et al. constructed a mathematical model to investigate the enhanced antitumor efficacy observed with NHS-muIL12 and Avelumab combination therapy in preclinical cancer models [14,26]. In addition to analyzing the model, they found by using numerical simulations that this combination therapy requires only about one-third the individual drug doses for tumor control compared to monotherapy.

In their simulation studies they rigorously estimated parameter values from the literature and the remaining parameters were estimated by fitting the model to cancer treatment experiments that were conducted on mice in [14]. These experiments included: a) Isotype control (no drug); b) NHS-muIL12 (2 µg); c) NHS-muIL12 (10 µg); d) Avelumab (200 µg); e) Avelumab (200 µg) and NHS-muIL12 (2 µg) and f)Avelumab (200 µg) and NHS-muIL12 (10 µg). BALB/c mice bearing orthotopic EMT-6 tumors ( 100 mm^3^) were treated with Avelumab on days 0, 3, and 6, while NHS-muIL12 was administered as a single dose on day 0. Model fitting was systematic in the sense that they sequentially fit the model to data using more complexity as more treatments were introduced. Specifically, they used the no drug case to estimate the proliferation rate of the tumor cells (*r*) and the kill rate of tumor cells by T cells (*η*). From there, they estimated *A*_2_ by fitting the model to the NHS-muIL12 (2 µg) data and similarly they estimated 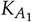 using the Avelumab (200 µg) data.

While the model by Nikolopoulou et al. provides a valuable framework, it fails to recapitulate the non-monotonic dynamics observed in the low-dose combination therapy. In this study, we posit that this discrepancy arises from the model’s assumption of a constant PD-L1 tumor expression propensity, *ϵ*. We present an iterative model refinement, resulting in a model with a dynamic *ϵ*, that successfully explains these complex dynamics. This work provides a quantitative framework for understanding adaptive immune resistance and establishes a platform for developing novel, model-derived biomarkers to predict therapeutic outcomes. Figure 2 shows the model with treatment-dependent *ϵ*, illustrating that allowing *ϵ* to vary across treatments both improves the model fit and captures tumor adaptation to the immune response under different therapies.

**Figure 2.**
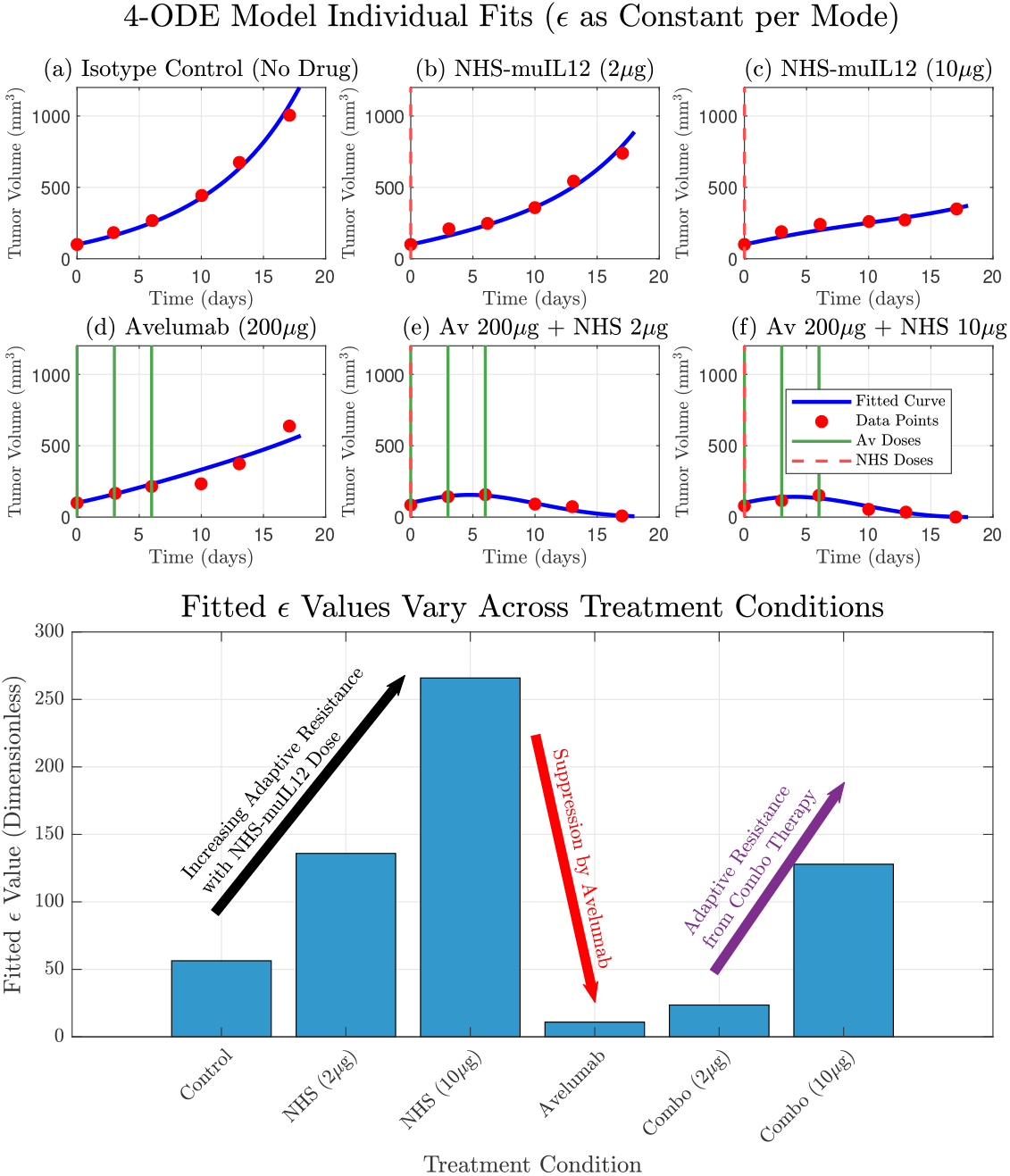
The original model by Nikolopoulou et al. adapted with therapy-specific PD-L1 expression (*ϵ*) accurately recapitulates experimental data and reveals the dynamics of adaptive resistance. (Top) The model’s simulated tumor volume (blue curves) shows a good fit to the experimental data (red circles) for all six treatment conditions. This fit was achieved by treating *ϵ* as a constant parameter that was individually fitted for each specific therapy. (Bottom) The resulting fitted values for *ϵ* are displayed for each condition. The values show a clear dose-dependent upregulation of *ϵ* in response to NHS-muIL12 monotherapy, a hallmark of adaptive resistance. Conversely, therapies including Avelumab show a strong suppression of the effective *ϵ* value, with a slight increase in the combination therapies due to the presence of NHS-muIL12.

**Figure 3.**
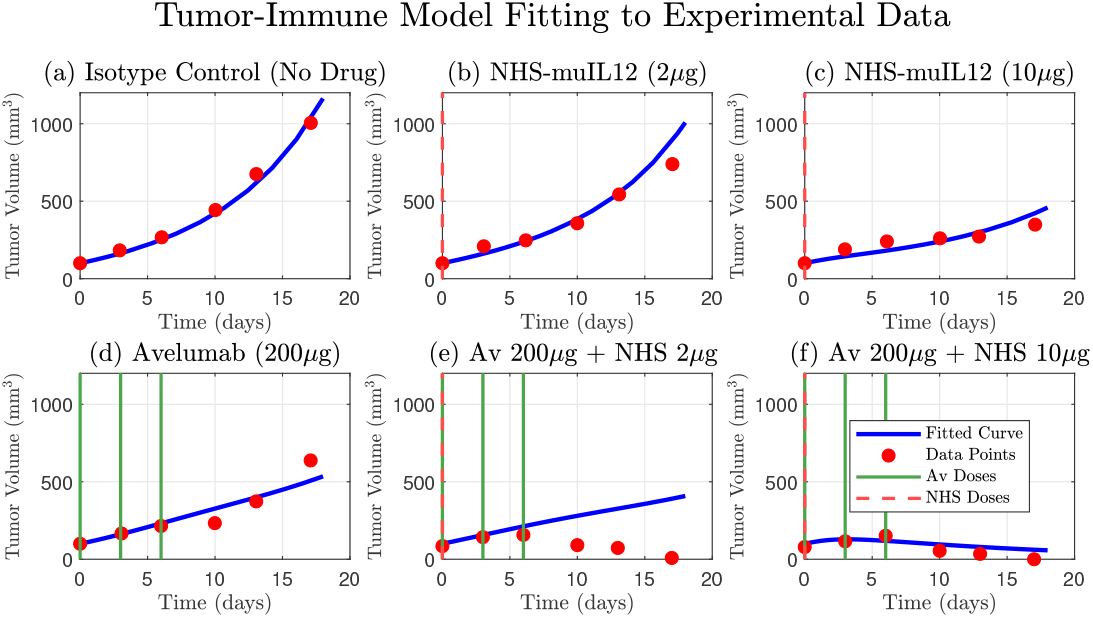
Comparison of the original 4-ODE model simulation against experimental tumor growth data. The model, parameterized as in Nikolopoulou et al. [26], accurately describes the tumor growth kinetics for the isotype control group (a) and the partial efficacy of the NHS-muIL12 (b, c) and Avelumab (d) monotherapies. The simulated tumor volume (blue curve) is shown against the experimental data (red circles). The model also captures the strong synergistic effect and tumor regression observed in the high-dose combination therapy (f). However, a key finding is the model’s inability to reproduce the tumor dynamics seen in the low-dose combination therapy data (e), where the tumor volume initially increases before decreasing. This discrepancy highlights a limitation in the original model formulation and suggests that a key biological mechanism is not being accounted for. Dosing schedules for Avelumab and NHS-muIL12 are indicated by solid green and dashed red lines, respectively.

## 2. Materials and Methods

### 2.1. Formulation of the Mathematical Model

We introduce the cancer treatment model first proposed by Nikolopoulou et al., who considered the micro tumor environment consisting of tumor cells and activated T-cells. Their mathematical model portrayed in Quick Guide 1 (Box 2.1) describes the interaction between tumor cells, activated T-cells, the anti–PD-L1 antibody Avelumab, and the immunostimulant NHS-muIL12. What follows is a narrative summary of the equations and assumptions.

In this model, tumor cells (mm^3^) grow exponentially but are counteracted by T-cell-mediated killing. T-cell population (mm^3^) increases in response to both tumor antigen stimulation and the immune-boosting effects of NHS-muIL12, while also undergoing natural turnover. The two drugs, Avelumab and NHS-muIL12, are modeled through their pharmacokinetics, with infusion inputs (*γ*_*i*_(*t*)) and clearance terms 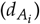.

The key features of the model are the mechanistic approaches to immune stimulation and immune evasion. T-cells are produced at a basal rate and stimulated by the presence of NHS-muIL12 in the micro tumor environment. The stimulation rate is assumed to be proportional to the T-cell population and NHS-muIL12 dosage whereas basal production is assumed constant. To incorporate immune evasion, the model temporally tracks the amount of PD-1/PD-L1 complex, *Q*, which is assumed to be generated from T-cells and tumor cells, but decreases with the anti PD-L1 agent Avelumab. In particular, the model assumes that PD-1 is expressed on the surface of T-cells and PD-L1 is expressed by both T-cell and tumor cell surfaces. We are interested in the latter because this enables the tumors ability to evolve and evade the immune system. Detailed model formulation and explanations can be found in [26] and are summarized in Quick Guide 1 (Box 2.1) below.

### 2.2. Experimental Data

Tumor volume data were obtained from digitizing the data from Figure 1B of Xu et al. using PlotDigitizer (https://plotdigitizer.com/app) [14,27]. However, we leave out the outlier data found in Avelumab (200 µg) and NHS-muIL12 (10 µg) combination case. Xu et al. rigorously investigated the antitumor efficacy of NHS-muIL12 and Avelumab, both as single agents and in combination, across two distinct preclinical cancer models with the goal of determining whether combination therapy with NHS-muIL12 and the anti-PD-L1 antibody Avelumab can enhance antitumor efficacy in preclinical models relative to monotherapies [14]. In particular, to generate the EMT-6 tumor data, BALB/c mice were inoculated with 0.5 × 10^6^ EMT-6 tumor cells orthotopically in the mammary fat pad. Mice were randomized into treatment groups when tumors reached the desired volume (day0) and treatment was initiated on day 0 [14]. Avelumab or isotype control were injected intravenously on days 0, 3, and 6 for EMT-6 tumor bearing mice. NHS-muIL12 was injected as a single subcutaneous dose on day 0 [14].

### 2.3. Updated Parameter Values

To understand tumor evolution and adaptation, we take the original model (see Box 2.1) and fit *ϵ* across the different therapies. In this way, we can understand the tumors adaptive response as different treatments are used.

We use updated parameter values either found from literature or values that were initially derived by Nikolopoulou et. al. but were not used. In particular, we take *λ*_T8*I*12_ = 4.15 day^−1^ as was initially used by Lai and Friedman to account for CD8^+^ T cells [28]. In addition, we update the degradation rates for Avelumab (*A*_1_) and NHS-muIL12 (*A*_2_), since the original parameters were derived from human data, and the experiments by Xu et. al. used mice. Murine-specific literature provides the following more accurate values:

- Avelumab: half-life *≈*44.6 hours = 1.86 days = *⇒* 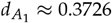 day^−1^[29].
- NHS-muIL12: half-life *≈* 9.5 days = *⇒* 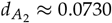 day^−1^[30].

#### Quick Guide 1

**Original Model Equations and Assumptions**

Here we summarize the model from Nikolopoulou et al. [26] which we will adapt for this study. The model simplifies the tumor microenvironment to consist of two primary interacting cell populations: tumor cells, with volume *V*(*t*) (mm^3^), and effector T-cells, with volume *T*(*t*) (mm^3^). The model also tracks the concentration of the two therapeutic agents: the anti-PD-L1 antibody Avelumab, *A*_1_(*t*), and the immunocytokine NHS-muIL12, *A*_2_(*t*). The dynamics of these populations and agents are governed by the following system of ordinary differential equations:

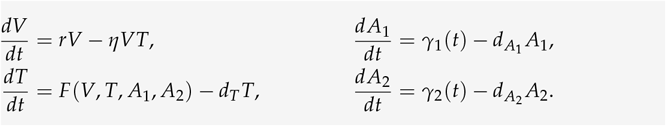

The top equation in the left column describes tumor growth, which is assumed to grow at an exponential rate (intrinsic growth rate *r*), but is reduced by T-cell-mediated killing at a rate *η* based on the law of mass action. The bottom left equation models the rate of change of the T-cell population, where the function *F* represents stimulation by the tumor and drug treatments, and decays at a rate *d*_*T*_. Finally, the pharmacokinetics of the anti-PD-L1 antibody Avelumab (*A*_1_) and the immunostimulant NHS-muIL12 (*A*_2_) are modeled, where *γ*_*i*_(*t*) terms represent drug administration and 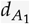 and 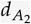 are the respective clearance rates.

The T-cell activation function *F* quantifies the production and stimulation of the immune system and is given by:

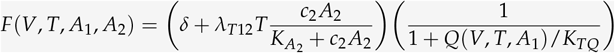

where *Q*(*V, T, A*_1_) quantifies the effective inhibition via the PD-1/PD-L1 complex. The first term in the parentheses represents the basal production and stimulatory component. In particular, *F* increases with increasing values of the immunostimulate *A*_2_ which is assumed to follow Michaelis-Menten dynamics and is proportional to *T*. Furthermore, it is assumed that there is a background production at rate *δ*. The second term in the product represents the immunosuppressive effect of the PD-1/PD-L1 immune checkpoint, and how it is reversed by the checkpoint inhibitor Avelumab (*A*_1_), through the PD-1/PD-L1 complex, *Q*. As *Q* increases, this second term gets smaller and hence reduces the value for *F. Q* is a function dependent on *V, T* and *A*_1_ and has the following form given by:

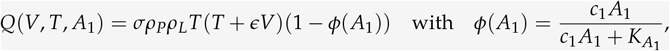

where *σ* represents the fraction of complex association and dissociation which was derived in [26]. It assumes that the complex *Q* increases proportional to *T* and *V*, but decreases with more anti-PD-L1 antibody *A*_1_ through the term 1 −*ϕ* which assumes *ϕ* is a Michaelis-Menten response curve. *ρ*_*P*_ and *ρ*_*L*_ are the expression levels of PD-1 and PD-L1, respectively. In this model, *ϵ* represents the effective level of tumor-derived PD-L1 expressed on the tumor cell surface, serving as a proxy for the strength of PD-L1-related immune suppression rather than a direct molecular count. For a more detailed description of the model and parameter meanings see [26] and Table A1.

Lastly, we used the original values 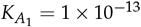 and 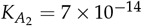 that were derived from literature values in [26]. We summarize the new parameter list in Table A1.

### 2.4. Model Refinement: Drug- and Tumor-Size–Dependent ϵ

While some tumors may have a baseline level of PD-L1 expression, it is not a fixed property. The tumor’s upregulation of PD-L1 is a defensive counter-adaptation. By increasing PD-L1 expression, the tumor can evade the very T-cells that were activated to attack it, creating a negative feedback loop that suppresses the immune response. A constant *ϵ* would completely ignore this crucial dynamic feedback, leading to an oversimplified and potentially inaccurate representation of the system’s dynamics, particularly for combination therapies. To that end we develop a differential equation for *ϵ* that depends on tumor volume *V*, drug treatments *A*_1_, *A*_2_ and itself, *ϵ*. We again provide a narrative description of the equation below. For a more detailed explanation of the derivation for this governing equation we point interested readers to Quick Guide 2 (Box 2.4).

We assume that dynamic PD-L1 expression increases at a rate that is proportional to tumor size and saturates at a maximum volume [31,32]. In addition, PD-L1 naturally decays over time, and drug-mediated degradation or suppression by Avelumab can further reduce its levels. We assume that the former follows an exponential decay while the latter follows a saturation function given by a Michaelis-Menten equation. Building in our assumption from the model where *ϵ* was held constant (see Figure 2), we further assume that *ϵ* increases with NHS-muIL12, we assume that PD-L1 expression increases as an adaptive response by tumor cells to immunostimulants.

### 2.5. Parameter Estimation

Model parameters were estimated by fitting the models to the experimental tumor volume data using the lsqnonlin function in MATLAB, which minimized the sum of squared errors (SSE) between the model simulation and the data. This process was performed for both models. For the base model, the PD-L1 expression parameter, *ϵ*, was fitted as a unique constant for each of the six therapies individually. For the final model the five key parameters governing the dynamics of *ϵ* from its own differential equation (Equation 1) were optimized simultaneously across all six experimental datasets.

## 3. Results

### 3.1. Base Model Limitations Highlight the Need for Dynamic ϵ

To empirically investigate the context-dependent nature of tumor PD-L1 expression, we performed individual fits of the core model where *ϵ* was treated as a constant parameter, fitted independently for each of the six treatment therapies. This approach allowed us to understand the tumor’s adaptive response as different treatments were used. The results from these individual fits demonstrated that the optimal *ϵ* value varied across treatments. For instance, NHS-muIL12 monotherapy (therapies 2 and 3) led to a substantial increase in the fitted *ϵ* compared to the Isotype Control (therapy 1), suggesting a tumor counter-adaptation by upregulating PD-L1 in response to enhanced immune stimulation. Conversely, Avelumab monotherapy (therapy 4) resulted in a markedly lower fitted *ϵ*, indicating a suppression of the tumor’s effective PD-L1 expression. These empirical findings provide strong motivation for the subsequent development of more mechanistic representations for *ϵ*. Model fits are presented in Figure 2 and we show how *ϵ* varies across therapies.

#### Quick Guide 2

**Dynamic PD-L1 Incoporation**

To account for the adaptive nature of the tumor’s PD-L1 expression, we introduced a fifth ordinary differential equation to govern the dynamics of the state variable *ϵ*. The rate of change of *ϵ* is modeled as a balance between production and degradation forces:

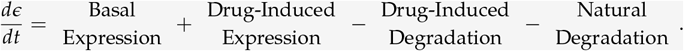

We assume that tumors possess a basal production rate of PD-L1 expression that is independent of external stimulation. Here, that rate follows Michaelis-Menten dynamics, where the production rate increases with tumor size but approaches a maximum:

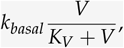

here *k*_*basal*_> represents the maximum production rate of PD-L1 from tumor cells and *K*_*V*_ is the half-saturation constant.

A fundamental mechanism of tumor immune escape is the upregulation of PD-L1 in response to an active anti-tumor immune attack. This process is primarily driven by pro-inflammatory cytokines, most notably interferon-gamma (IFN-*γ*), which is secreted by activated T-cells upon tumor antigen recognition. The PD-L1 production rate is modeled as being stimulated by NHS-muIL12 (*A*_2_) in a tumor size-dependent manner, representing IFN-*γ*-mediated upregulation. This formulation captures adaptive immune resistance, in which immune-activating signals from the drug induce PD-L1 expression as a tumor defense mechanism. Based on this discussion, we assume functional forms that follow a saturation curve for both *A*_2_ and *V*:

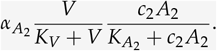

where *K*_*ϵ*_ and 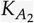 are the half-saturation constants for *V* and *A*_2_, respectively and 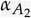 represents the maximum rate of PD-L1 production by NHS-muIL12 stimulation.

Avelumab is a monoclonal antibody that specifically binds to the PD-L1 protein on cancer cells. By doing so, it prevents PD-L1 from attaching to the PD-1 receptor on T-cells, which would normally send an inhibitory signal to deactivate them. This blockade allows the T-cells to remain active and effectively target and destroy the tumor cells. To this end, we assume the drug-mediated suppression of PD-L1 by Avelumab (*A*_1_) is given by the following:

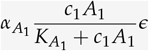

where *c*_1_ and 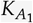 are defined as before and 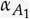 represents the maximum rate of drug-mediated suppression of PD-L1 by Avelumab.

Finally, we assume that tumor-derived PD-L1 expression decays at rate *d*_*ϵ*_. The governing equation for the change in tumor PD-L1 expression level (*ϵ*) over time is

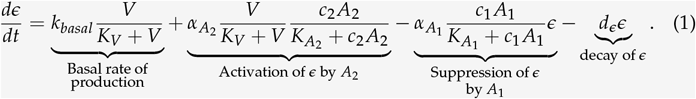

Since treatment pulses are introduced at *t* = 0 in the simulation framework, the initial value of *ϵ* was computed without drug intervention. We further assumed that before treatment, *ϵ* reaches its quasi–steady state determined by the tumor volume at time zero. This leads to

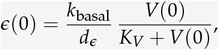

with *V*(0) = 100 representing the tumor volume at the start of the simulation as taken by Nikolopoulou et al., which aligns with experimental observations [14,26].

### 3.2. Dynamic ϵ Improves Fit and Explanatory Power

The model that included the dynamic *ϵ* (Equation 1) successfully fit the full spectrum of experimental outcomes, including the monotherapy responses and the synergistic tumor regression in the low and high-dose combination therapy Figure 4. The model’s ability to accurately capture all six datasets with a single set of parameters demonstrates its robustness and explanatory power. Estimated parameters are shown in Table 1.

**Table 1.** Additional parameter values used in the dynamic *ϵ* model. All parameters have units time^−1^.

| Var. | Meaning | Fitted Value |
| --- | --- | --- |
| $k_{\text{basal}}$ | Basal proliferation rate | 30.5441 |
| $K_V$ | PD-L1 Upregulation Sensitivity | 50 |
| $\alpha_{A_1}$ | Decay rate induced by Avelumab | 38.2653 |
| $\alpha_{A_2}$ | Proliferation induced by NHS-muIL12 | 643.6397 |
| $d_\epsilon$ | Decay rate | 0.4409 |

**Figure 4.**
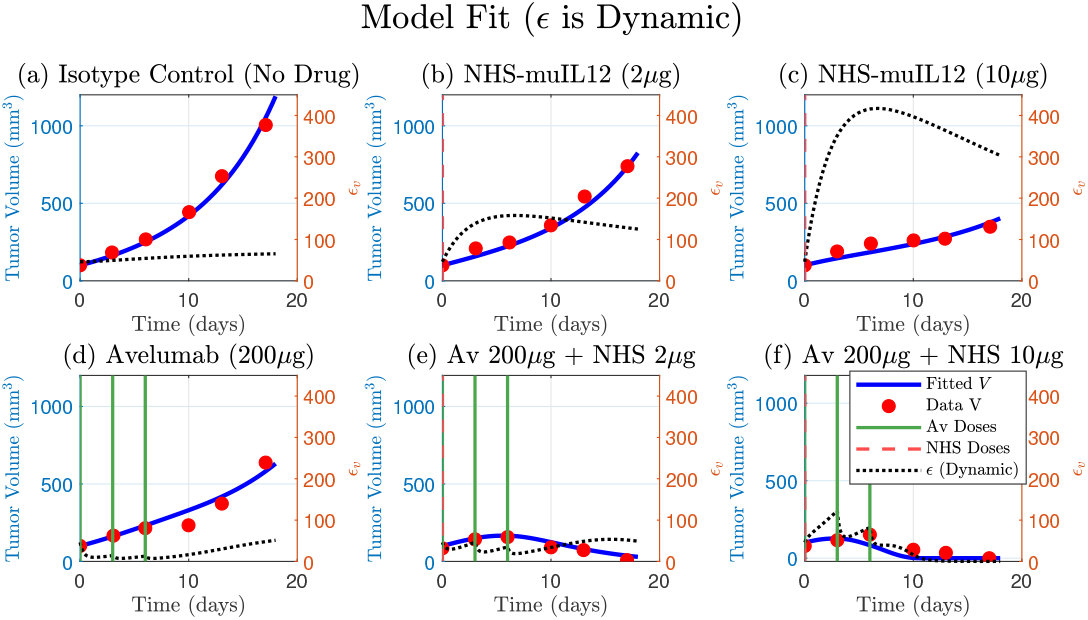
A single set of fitted parameters for the model that includes a dynamic PD-L1 expression (Equation 1) successfully fits the full spectrum of therapeutic outcomes. The model’s simulated tumor volume (blue curve) is plotted against tumor data (red circles) for all six treatments. The model accurately captures the monotherapy responses (panels b, c and d), the tumor regression in the low and high-dose combination therapies. The corresponding simulated trajectory *ϵ* (dashed-black, right y-axis) provides a mechanistic basis for these responses, qualitatively demonstrating adaptive resistance by the tumor. Avelumab and NHS-muIL12 doses are indicated by solid green and dashed vertical lines, respectively. Four parameters were fitted: *α*_*basal*_, 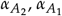 and *d*_*ϵ*_ (see Table 1 for values).

Furthermore, the simulated trajectory of *ϵ* itself provides a mechanistic explanation for the observed tumor dynamics. In the NHS-muIL12 monotherapies, the model predicts an early sharp and sustained increase in *ϵ*, quantitatively simulating the process of adaptive immune resistance. Conversely, in the presence of Avelumab, the model shows a strong suppression of *ϵ*. This alignment between the model’s internal dynamics and the known biological mechanisms validates the model’s structure and confirms that the dynamic regulation of PD-L1 is a key determinant of the therapeutic outcome.

### 3.3. Model Comparisons (RSS and AIC)

We compared two alternative models: the model with a dynamic *ϵ* and a constant *ϵ* model, in which *ϵ* was fitted independently for each treatment. In the constant epsilon model, *ϵ* was fitted independently for each of the six experimental conditions. Although this represents the same biological parameter, it was treated as six free parameters in the AIC calculation, since each condition was assigned its own fitted value. This approach provides a fair comparison with the dynamic *ϵ* model, where *ϵ* is estimated globally with Equation 1 with five parameters.

We summarize residual sum of squares (RSS) and Akaike information criterion (AIC) here, but provide detailed results in Table 2. Both approaches supported the central hypothesis that *ϵ* is not fixed but instead varies across treatment conditions due to tumor evolution and adaption. At the global level, the constant *ϵ* model provided a better fit, yielding a total RSS of 3.52 × 10^4^ and an AIC of 259.8, compared to an RSS of 4.76 × 10^4^ and AIC of 268.8 for the dynamic *ϵ* model. These results indicate that the constant *ϵ* formulation better explains the data set overall. However, our main result still holds that *ϵ* is dynamically changing across treatments since both models incorporate this hypothesis.

**Table 2.**
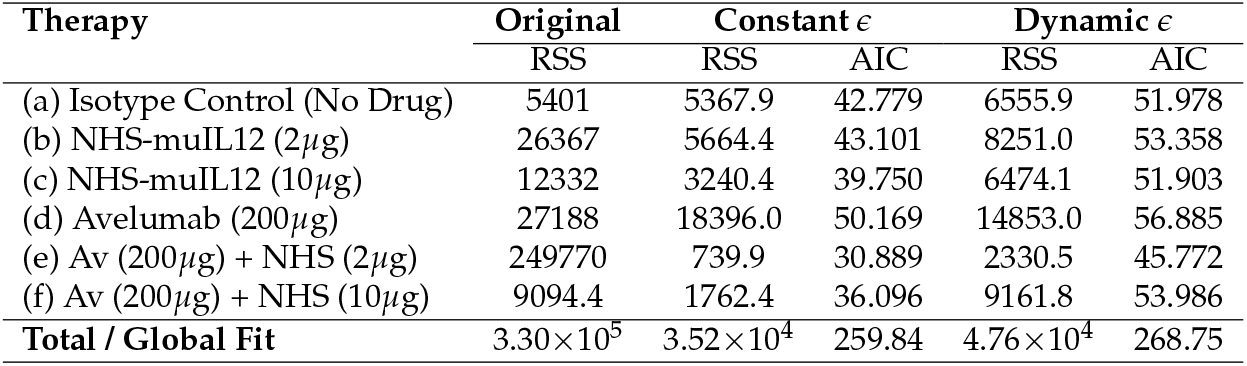
Model comparison between the original, constant *ϵ*, and dynamic *ϵ* model formulations across experimental conditions. Residual sum of squares (RSS) and Akaike information criterion (AIC) values are shown. Lower values indicate a better fit. In this comparison, the original model uses the updated parameters as discussed in Section 2.3

When broken down by treatment, the constant *ϵ* model generally achieved lower RSS and AIC values (Table 2). For example, in the low dose combination treatment, the residual error decreased substantially under the constant *ϵ* model (739.9 vs. 2330.5). Similarly, for the high dose combination therapy the constant *ϵ* model reduced RSS by more than half (3240.4 vs. 6474.1). The one notable exception was the Avelumab-only treatment, where the dynamic *ϵ* model produced a better fit (RSS = 14,853 vs. 18,396).

Overall, these results confirm that both modeling approaches are consistent with the hypothesis of a changing *ϵ*, while the constant *ϵ* model provides a more accurate description across the full set of experimental conditions.

## 4. Discussion

### 4.1. Summary of Key Findings

In this study, we developed a mechanistic model that qualitatively explains the complex dynamics of combination therapy first considered by Nikolopoulou et al., that was derived from experimental data [14]. We observed that *ϵ* was a dynamically changing parameter rather than a constant by fitting it individually across all therapies (Figure 2, bottom panel). This intermediate step improved the fits and provided strong quantitative evidence that the tumor was actively adapting its PD-L1 expression in response to the different therapies. However, this descriptive approach lacked predictive power and motivated the development of a dynamic representation of PD-L1 expression, where *ϵ* is governed by its own mechanistic differential equation. This provided a balance between predictive power while still improving model fits to data, in particular the non-monotone dynamics found in the low-dose combination therapy, see Figures 2, 3 and 4. Ultimately, both the dynamic and constant *ϵ* models supported the hypothesis that *ϵ* varies across therapies. While the constant epsilon model provided a better overall fit (lower RSS and AIC), the dynamic epsilon model captured treatment-specific effects such as Avelumab monotherapy.

### 4.2. Adaptive Resistance via PD-L1 Upregulation

The final model provides a quantitative validation for the biological mechanism of adaptive immune resistance. Our simulations of NHS-muIL12 monotherapy accurately predict a dramatic increase in *ϵ*, leading to T-cell suppression and subsequent treatment failure. This directly reflects the IFN-*γ*-dependent PD-L1 upregulation demonstrated experimentally by Fallon et al., who conclusively proved this link using IFN-*γ* knockout mice [33]. The differential equation governing *ϵ* in our model includes the parameter *α*_*NHS*_ which explicitly captures this process. We chose to link PD-L1 upregulation directly to the presence of NHS-muIL12 rather than explicitly modeling the intermediate IFN-*γ* step, as this simpler formulation is more robust and avoids further parameter identifiability issues while still capturing the essential dynamics.

Our finding that dynamic PD-L1 expression is a critical determinant of therapeutic outcome is complemented by the work of Lai and Yu [34]. Through a stability analysis of a similar model that included T-cell exhaustion, they independently identified tumor PD-L1 expression as a sensitive parameter that governs the bistability of tumor-free and tumorous states, reinforcing its central role in mediating immune escape.

### 4.3. Alternative Mechanisms and Model Limitations

While our model mechanistically attributes treatment failure to the upregulation of the tumor’s PD-L1 expression, another biologically plausible mechanism that could explain the observed relapse is T-cell exhaustion in the tumor microenvironment [35,36]. This phenomenon describes a state of T-cell dysfunction that arises from chronic exposure to tumor antigens. Over time, persistent stimulation can lead to a progressive loss of effector functions, such as the ability to proliferate and secrete cytotoxic molecules, even if the PD-1/PD-L1 blockade is maintained [35]. In the context of our model, this would mean that even with a dynamically changing PD-L1 expression (*ϵ*), the T-cell population (*T*) itself would become less effective at killing tumor cells. Future iterations of this model could incorporate T-cell exhaustion by making the T-cell death rate (*d*_*T*_) a function of a proxy for cumulative antigen exposure, or by adding a separate, “exhausted” T-cell population. Distinguishing the relative contributions of adaptive resistance via PD-L1 upregulation versus intrinsic T-cell exhaustion remains a key challenge and an important direction for future investigation.

An alternative interpretation of our model’s dynamic *ϵ* is that it represents the consequence of clonal selection within a heterogeneous tumor [37]. While the biological reality is likely a continuous spectrum of resistant cells, it is simpler to conceptualize this as a mixture of two populations: therapy-sensitive cells (with a low potential for PD-L1 expression) and a pre-existing sub-clone of intrinsically resistant cells (with a high capacity for PD-L1 upregulation) [38–43]. The immunotherapy then acts as a strong selective pressure, efficiently eliminating the sensitive population at one end of the spectrum. This allows the more resistant clones, which were initially a small fraction of the tumor, to survive and proliferate, eventually shifting the entire population’s distribution towards higher resistance. As this evolutionary process occurs, the average PD-L1 expression potential of the whole tumor increases, which is precisely what our model captures through the dynamic increase of the single epsilon parameter under NHS-muIL12 therapy (Figure 2). Therefore, our model’s framework can be seen as an effective representation of the bulk tumor dynamics that result from this underlying process of clonal evolution across a resistance spectrum.

## 5. Conclusions

This study demonstrates the power of an iterative mathematical modeling approach to quantitatively dissect the mechanisms of adaptive resistance in combination immunotherapy. By showing that incorporating therapy-dependent, nonconstant regulation of PD-L1 enabled a biologically grounded mechanism to reproduce experimental observations, we formulated PD-L1 tumor expression as a dynamic variable (*ϵ*), thereby providing a mechanistic basis for both therapeutic synergy and treatment failure within the original model. This work builds onto a robust in silico platform that can be leveraged to design and test novel therapeutic strategies to overcome the challenge of tumor immune escape.

## Author Contributions

Conceptualization, B.P., D.D., A. Kalizhanova, A. Kashkynbayev, Y.K.; methodology, B.P., A. Kalizhanova, Y.K.; software, B.P., A. Kalizhanova; validation, B.P., D.D., A. Kalizhanova; formal analysis, B.P.; investigation, A. Kalizhanova, A. Kashkynbayev, A. T., B.P., D.D., Y.K.; resources, A. Kashkynbayev, B.P., Y.K.; data curation, B.P., D.D.; writing—original draft preparation, B.P.; writing—review and editing, A. Kalizhanova, A. Kashkynbayev, A. T., B.P., D.D., Y.K.; visualization, A. T., B.P.; supervision, A. Kashkynbayev, Y.K.; project administration, B.P., Y.K.; funding acquisition, A. Kashkynbayev, B.P., Y.K. All authors have read and agreed to the published version of the manuscript.

## Appendix A

Here we provide the parameter values that were used for the model fittings.

**Table A1.** Model parameters, their meanings, values, and references.

| Var. | Meaning | Value | Ref. |
| --- | --- | --- | --- |
| $r$ | Tumor cell growth rate | $0.213 \text{ day}^{-1}$ | [26] |
| $\eta$ | Kill rate of tumor cells by T cells | $1 \text{ mm}^{-3} \text{ day}^{-1}$ | [26] |
| $\delta$ | Source of T cell activation | $0.02 \text{ mm}^3/\text{day}$ | [26] |
| $\lambda_{T_{IL12}}$ | Activation rate of T cells by IL-12 | $4.15 \text{ day}^{-1}$ | [28] |
| $K_{TQ}$ | Inhibition of T cells by PD-1–PD-L1 | $1 \times 10^{-13} \text{ mm}^6$ | [26] |
| $K_{A_1}$ | Dissociation constant of PD-L1–anti-PD-L1 | $1 \times 10^{-13} \text{ mol/L}$ | [26] |
| $K_{A_2}$ | Dissociation constant of PD-L1–anti-PD-L1 | $1 \times 7 \times 10^{-14} \text{ mol/L}$ | [26] |
| $d_T$ | Death rate of T cells | $0 - 0.5 \text{ day}^{-1}$ | [44] |
| $d_{A_1}$ | Degradation rate of Anti-PD-L1 | $0.3726 \text{ day}^{-1}$ | [29] |
| $d_{A_2}$ | Degradation rate of NHS-muIL12 | $0.0730 \text{ day}^{-1}$ | [30] |
| $\rho_p$ | Expression level of PD-1 | $3.19 \times 10^{-7} - 8.49 \times 10^{-7}$ | [28] |
| $\rho_L$ | Expression level of PD-L1 | $3.56 \times 10^{-7} - 1.967 \times 10^{-6}$ | [28] |
| $\epsilon$ | PD-L1 expression from tumor | – | fitted |
| $\sigma$ | Complex assoc./dissoc. fraction | $0.01 \text{ mm}^{-3}$ | [26] |
| $\gamma_1$ | Infusion rate of Avelumab | $1 \times 10^{-7} - 9 \times 10^{-5} \text{ g/day}$ | [26] |
| $\gamma_2$ | Infusion rate of NHS-muIL12 | $1 \times 10^{-9} - 2 \times 10^{-6} \text{ g/day}$ | [26] |
| $c_1$ | Conversion constant for $A_1$ | $55^{-1} \times 10^{-7} - 55^{-1} \times 10^{-6}$ | [26] |
| $c_2$ | Conversion constant for $A_2$ | $75^{-1} \times 10^{-7} - 75^{-1} \times 10^{-6}$ | [26] |

## Disclaimer/Publisher’s Note

The statements, opinions and data contained in all publications are solely those of the individual author(s) and contributor(s) and not of MDPI and/or the editor(s). MDPI and/or the editor(s) disclaim responsibility for any injury to people or property resulting from any ideas, methods, instructions or products referred to in the content.

## References

1. Burrell, R.A.; McGranahan, N.; Bartek, J.; Swanton, C. The causes and consequences of genetic heterogeneity in cancer evolution. Nature 2013, 501, 338–345. Publisher: Nature Publishing Group, 10.1038/nature12625.

2. Gerlinger, M.; McGranahan, N.; Dewhurst, S.M.; Burrell, R.A.; Tomlinson, I.; Swanton, C. Cancer: Evolution Within a Lifetime. Annual Review of Genetics 2014, 48, 215–236. Publisher: Annual Reviews, 10.1146/annurev-genet-120213-092314.

3. Vendramin, R.; Litchfield, K.; Swanton, C. Cancer evolution: Darwin and beyond. The EMBO Journal 2021, 40, e108389. Publisher: John Wiley & Sons, Ltd, 10.15252/embj.2021108389.

4. Pardoll, D.M. The blockade of immune checkpoints in cancer immunotherapy. Nature Reviews Cancer 2012, 12, 252–264. Publisher: Nature Publishing Group, 10.1038/nrc3239.

5. Schilsky, R.L. Implementing personalized cancer care. Nature Reviews Clinical Oncology 2014, 11, 432–438. Publisher: Nature Publishing Group, 10.1038/nrclinonc.2014.54.

6. Lin, H.H.; Wei, N.C.; Chou, T.Y.; Lin, C.C.; Lan, Y.T.; Chang, S.C.; Wang, H.S.; Yang, S.H.; Chen, W.S.; Lin, T.C.; et al. Building personalized treatment plans for early-stage colorectal cancer patients. Oncotarget 2017, 8, 13805–13817. 10.18632/oncotarget.14638.

7. Han, Y.; Liu, D.; Li, L. PD-1/PD-L1 pathway: current researches in cancer. American Journal of Cancer Research 2020, 10, 727–742.

8. Juneja, V.R.; McGuire, K.A.; Manguso, R.T.; LaFleur, M.W.; Collins, N.; Haining, W.N.; Freeman, G.J.; Sharpe, A.H. PD-L1 on tumor cells is sufficient for immune evasion in immunogenic tumors and inhibits CD8 T cell cytotoxicity. The Journal of Experimental Medicine 2017, 214, 895–904. 10.1084/jem.20160801.

9. Collins, J.M.; Gulley, J.L. Product review: avelumab, an anti-PD-L1 antibody. Human Vaccines & Immunotherapeutics 2019, 15, 891–908. Publisher: Taylor & Francis _eprint: https://doi.org/10.1080/21645515.2018.1551671, 10.1080/21645515.2018.1551671.

10. Vaishampayan, U.; Schöffski, P.; Ravaud, A.; Borel, C.; Peguero, J.; Chaves, J.; Morris, J.C.; Kotecki, N.; Smakal, M.; Zhou, D.; et al. Avelumab monotherapy as first-line or second-line treatment in patients with metastatic renal cell carcinoma: phase Ib results from the JAVELIN Solid Tumor trial. Journal for ImmunoTherapy of Cancer 2019, 7, 275. 10.1186/s40425-019-0746-2.

11. Liu, K.; Tan, S.; Chai, Y.; Chen, D.; Song, H.; Zhang, C.W.H.; Shi, Y.; Liu, J.; Tan, W.; Lyu, J.; et al. Structural basis of anti-PD-L1 monoclonal antibody avelumab for tumor therapy. Cell Research 2017, 27, 151–153. Publisher: Nature Publishing Group, 10.1038/cr.2016.102.

12. Fallon, J.; Tighe, R.; Kradjian, G.; Guzman, W.; Bernhardt, A.; Neuteboom, B.; Lan, Y.; Sabzevari, H.; Schlom, J.; Greiner, J.W. The immunocytokine NHS-IL12 as a potential cancer therapeutic. Oncotarget 2014, 5, 1869–1884. 10.18632/oncotarget.1853.

13. Greiner, J.W.; Morillon II, Y.M.; Schlom, J. NHS-IL12, a Tumor-Targeting Immunocytokine. ImmunoTargets and Therapy 2021, 10, 155–169. Publisher: Dove Medical Press _eprint: https://www.tandfonline.com/doi/pdf/10.2147/ITT.S306150, 10.2147/ITT.S306150.

14. Xu, C.; Zhang, Y.; Rolfe, P.A.; Hernández, V.M.; Guzman, W.; Kradjian, G.; Marelli, B.; Qin, G.; Qi, J.; Wang, H.; et al. Combination Therapy with NHS-muIL12 and Avelumab (anti-PD-L1) Enhances Antitumor Efficacy in Preclinical Cancer Models. Clinical Cancer Research 2017, 23, 5869–5880. 10.1158/1078-0432.CCR-17-0483.

15. Altrock, P.M.; Liu, L.L.; Michor, F. The mathematics of cancer: integrating quantitative models. Nature Reviews Cancer 2015, 15, 730–745. Publisher: Nature Publishing Group, 10.1038/nrc4029.

16. Phan, T.; Crook, S.M.; Bryce, A.H.; Maley, C.C.; Kostelich, E.J.; Kuang, Y. Review: Mathematical Modeling of Prostate Cancer and Clinical Application. Applied Sciences 2020, 10, 2721. Publisher: Multidisciplinary Digital Publishing Institute, 10.3390/app10082721.

17. Brady, R.; Enderling, H. Mathematical Models of Cancer: When to Predict Novel Therapies, and When Not to. Bulletin of Mathematical Biology 2019, 81, 3722–3731. 10.1007/s11538-019-00640-x.

18. Rauf, S.; Smirnova, A.; Chang, A.; Liu, Y.; Jiang, Y. Immunogenic Cell Death: the Key to Unlocking the Potential for Combined Radiation and Immunotherapy. bioRxiv 2025. 10.1101/2025.02.14.638342.

19. Kuang, Y.; Nagy, J.D.; Eikenberry, S.E. Introduction to Mathematical Oncology; Chapman & Hall/CRC: Boca Raton, FL, 2016.

20. Serre, R.; Benzekry, S.; Padovani, L.; Meille, C.; André, N.; Ciccolini, J.; Barlesi, F.; Muracciole, X.; Barbolosi, D. Mathematical Modeling of Cancer Immunotherapy and Its Synergy with Radiotherapy. Cancer Research 2016, 76, 4931–4940. 10.1158/0008-5472.CAN-15-3567.

21. Bräutigam, K. Optimization of chemotherapy regimens using mathematical programming. Computers & Industrial Engineering 2024, 191, 110078. 10.1016/j.cie.2024.110078.

22. Hahnfeldt, P.; Panigrahy, D.; Folkman, J.; Hlatky, L. Tumor development under angiogenic signaling: a dynamical theory of tumor growth, treatment response, and postvascular dormancy. Cancer Research1999, 59, 4770–4775.

23. Reckell, T.; Nguyen, K.; Phan, T.; Crook, S.; Kostelich, E.J.; Kuang, Y. Modeling the synergistic properties of drugs in hormonal treatment for prostate cancer. Journal of Theoretical Biology 2021, 514, 110570. 10.1016/j.jtbi.2020.110570.

24. Meade, W.; Weber, A.; Phan, T.; Hampston, E.; Resa, L.F.; Nagy, J.; Kuang, Y. High Accuracy Indicators of Androgen Suppression Therapy Failure for Prostate Cancer—A Modeling Study. Cancers 2022, 14, 4033. 10.3390/cancers14164033.

25. Phan, T.; Weber, A.; Bryce, A.H.; Kuang, Y. The prognostic value of androgen to PSA ratio in predictive modeling of prostate cancer. Medical Hypotheses 2023, 176, 111084. 10.1016/j.mehy.2023.111084.

26. Nikolopoulou, E.; Eikenberry, S.E.; Gevertz, J.L.; Kuang, Y. Mathematical modeling of an immune checkpoint inhibitor and its synergy with an immunostimulant. Discrete and Continuous Dynamical Systems - B 2021, 26, 2133–2159. 10.3934/dcdsb.2020138.

27. PlotDigitizer: Version 3.1.6. https://plotdigitizer.com, 2025.

28. Lai, X.; Friedman, A. Combination therapy of cancer with cancer vaccine and immune checkpoint inhibitors: A mathematical model. PloS One 2017, 12, e0178479. 10.1371/journal.pone.0178479.

29. Zalba, S.; Contreras-Sandoval, A.M.; Martisova, E.; Debets, R.; Smerdou, C.; Garrido, M.J. Quantification of Pharmacokinetic Profiles of PD-1/PD-L1 Antibodies by Validated ELISAs. Pharmaceutics 2020, 12, 595. 10.3390/pharmaceutics12060595.

30. Unverdorben, F.; Richter, F.; Hutt, M.; Seifert, O.; Malinge, P.; Fischer, N.; Kontermann, R.E. Pharmacokinetic properties of IgG and various Fc fusion proteins in mice. mAbs 2016, 8, 120–128. 10.1080/19420862.2015.1113360.

31. Wang, X.; Teng, F.; Kong, L.; Yu, J. PD-L1 expression in human cancers and its association with clinical outcomes. 9, 5023–5039. 10.2147/OTT.S105862.

32. Zhao, Y.; Shi, Z.; Xie, Y.; Li, N.; Chen, H.; Jin, M. The association between PD-1 / PD-L1 expression and clinicopathological features in sarcomatoid renal cell carcinoma. 47, 163–168. 10.1016/j.asjsur.2023.06.065.

33. Fallon, J.K.; Vandeveer, A.J.; Schlom, J.; Greiner, J.W. Enhanced antitumor effects by combining an IL-12/anti-DNA fusion protein with avelumab, an anti-PD-L1 antibody. Oncotarget 2017, 8, 20558–20571. 10.18632/oncotarget.16137.

34. Lai, X.; Yu, T. Modeling Combination Therapies and T Cell Exhaustion Dynamics in the Tumor Under Immune Checkpoint Blockade. Bulletin of Mathematical Biology 2025, 87, 128. 10.1007/s11538-025-01507-0.

35. Zhang, Z.; Liu, S.; Zhang, B.; Qiao, L.; Zhang, Y. T Cell Dysfunction and Exhaustion in Cancer. Frontiers in Cell and Developmental Biology 2020, 8. Publisher: Frontiers, 10.3389/fcell.2020.00017.

36. Pauken, K.E.; Wherry, E.J. Overcoming T cell exhaustion in infection and cancer. Trends in Immunology 2015, 36, 265–276. Publisher: Elsevier, 10.1016/j.it.2015.02.008.

37. Gniadek, T.J.; Li, Q.K.; Tully, E.; Chatterjee, S.; Nimmagadda, S.; Gabrielson, E. Heterogeneous expression of PD-L1 in pulmonary squamous cell carcinoma and adenocarcinoma: implications for assessment by small biopsy. Modern Pathology 2017, 30, 530–538. 10.1038/modpathol.2016.213.

38. Portz, T.; Kuang, Y.; Nagy, J.D. A clinical data validated mathematical model of prostate cancer growth under intermittent androgen suppression therapy. AIP Advances 2012, 2, 011002. 10.1063/1.3697848.

39. Portz, T.; Kuang, Y. A mathematical model for the immunotherapy of advanced prostate cancer. In BIOMAT 2012; WORLD SCIENTIFIC, 2013; pp. 70–85. 10.1142/9789814520829_0005.

40. Baez, J.; Kuang, Y. Mathematical Models of Androgen Resistance in Prostate Cancer Patients under Intermittent Androgen Suppression Therapy. Applied Sciences 2016, 6, 352. Publisher: Multidisciplinary Digital Publishing Institute, 10.3390/app6110352.

41. Phan, T.; He, C.; Martinez, A.; Kuang, Y.; Phan, T.; He, C.; Martinez, A.; Kuang, Y. Dynamics and implications of models for intermittent androgen suppression therapy. Mathematical Biosciences and Engineering 2019, 16, 187–204. Cc_license_type: cc_by Primary_atype: Mathematical Biosciences and Engineering Subject_term: Research article Subject_term_id: Research article, 10.3934/mbe.2019010.

42. Phan, T.; Nguyen, K.; Sharma, P.; Kuang, Y. The Impact of Intermittent Androgen Suppression Therapy in Prostate Cancer Modeling. Applied Sciences 2019, 9, 36. Publisher: Multidisciplinary Digital Publishing Institute, 10.3390/app9010036.

43. Everett, R.A.; Packer, A.M.; Kuang, Y. Can Mathematical Models Predict the Outcomes of Prostate Cancer Patients Undergoing Intermittent Androgen Deprivation Therapy? Biophysical Reviews and Letters 2014, 09, 173–191. Publisher: World Scientific Publishing Co., 10.1142/S1793048014300023.

44. Nikolopoulou, E.; Johnson, L.; Harris, D.; Nagy, J.; Stites, E.; Kuang, Y. Tumour-immune dynamics with an immune checkpoint inhibitor. Letters in Biomathematics 2018, 5, S137–S159. Publisher Copyright: © 2018, © 2018 The Author(s). Published by Informa UK Limited, trading as Taylor & Francis Group., 10.1080/23737867.2018.1440978.

